# Verbal Labels Facilitate Tactile Texture Discrimination in a Perceptual Learning Task

**DOI:** 10.1101/2021.02.09.430389

**Authors:** Ishita Arun, Leslee Lazar

## Abstract

The influence of language on perceptual processes, referred to as the Whorfian hypothesis, has been a contentious issue. Cross-linguistic research and lab-based experiments have shown that verbal labels can facilitate perceptual and discriminatory processes, mostly in visual and auditory modalities. Here, we investigated whether verbal labels improve performance in a tactile texture discrimination task using natural textures. We also explored whether the grammatical category of these verbal labels plays a role in discrimination ability. In our experiments, we asked the participants to discriminate between pairs of textures presented to the fingertip after a five-day training phase. During the training phase, the tactile textures and English pseudowords were co-presented consistently in the congruent (experimental) condition and inconsistently in the incongruent (control) condition, allowing them to form implicit associations only in the former condition. The pseudoword verbal labels belonged to two grammatical categories, verb-like and noun-like. We found an improvement in the texture discrimination ability only for the congruent condition, irrespective of the grammatical category.

## Introduction

The linguistic relativity hypothesis, proposed by Whorf, suggests that language serves as a tool for perception and other mental processes^1^. Although it has some detractors^2,3^, there is a wealth of empirical studies that show that people perceive and categorize sensory information differently, based on their native language^4–7^. Native Russian speakers were able to discriminate between two shades of blue faster and more accurately as their language has two different words for these shades; dark blue (siniy) and light blue (goluboy). On the other hand, English speakers could not do so as their language only has one word for both shades^4^. Similarly, comparing English and Berinmo speaking subjects, English speakers distinguished shades of blue and green stimuli as different, whereas Berinmo language speakers, whose language has no distinct words for these two colours, failed to do so^5^. These studies show that linguistic features informed the formation of categories in the continuous spectrum of sensory information. This category formation enabled people to better discriminate stimuli across the category boundary than within^8,9^.

The linguistic features that affect category formation also include pseudoword verbal labels without semantic meanings^10,11^; infants learn to categorize sounds into words by merely hearing them as labels presented alongside visual stimuli^12^. Similarly, when adult participants were trained to categorize images of aliens as approachable or avoidable based on certain physical features, the group with pseudoword names for the aliens categorized them faster than the unnamed group^10^. In another study, pseudoword adjectives were as effective as conventional words to categorize visual stimuli based on the properties of roundedness and pointedness^13^.

The effect of verbal labels on perception can manifest in multiple ways, like, improving discrimination, categorization, speed of perception, and even bringing to attention previously invisible stimuli^4,10,13–15^. If a verbal label precedes the stimuli in visual detection tasks, then the target stimuli’ detection is better^14–16^. The verbal labels can also affect the pre-attentive mechanisms of perception, shown in an ERP study of Greek and English-speaking participants performing a color discrimination task involving blue and green colors^17,18^. The Greek language has two words for two shades of blue, for which the English language only has one word. Greek-speaking participants showed an increase in the visual mismatch negativity, a marker of pre-attentive change detection for blue shades, unlike English speakers.

Neuroscientific studies with brain imaging and lesion studies show a functional connection between brain areas that process language and sensory perception^19–23^. A functional magnetic resonance imaging (fMRI) study showed that when participants perform a perceptual color discrimination task, the brain’s word-finding regions (left posterior superior temporal gyrus and the inferior parietal lobule) showed greater co-activation for easy-to-name colors than difficult-to-name colors^19^. Applying cathodal transcranial direct-current current stimulation (tDCS) over the language processing areas, Broca’s and Wernicke’s areas, also impaired perceptual categorization of visual stimuli^20,22^. Further evidence from studies on Aphasic patients with lesions in Wernicke’s area and loss of their linguistic abilities also showed impaired ability to categorize colors and facial expressions^21^.

The effects of verbal labels on perception have been well studied in the visual and auditory modalities^4,5,10,24–27^. Information on how the perceptual systems are affected by linguistic cues in other modalities are not yet well studied. Recently, a few studies addressed this issue in the olfactory and somatosensory systems^23,28,29^. Given the differences in the processing mechanisms and the nature of the stimulus features, it is imperative to understand the role of verbal labels in multiple sensory modalities to derive common principles of verbal labels’ effect on perception in general.

To this end, we studied the effect of verbal labels on discriminating natural textures using fingertips in an active touch paradigm. The tactile sensory experience is described by a rich linguistic vocabulary, especially in the perception of textures^30^. Texture is an essential attribute of a haptic object and refers to a material surface’s inherent microstructural properties, consisting of multiple dimensions such as width or spacing of microstructural surface patterns, thermal qualities, friction, etc^31–33^. People tend to report touching the textures using adjectives in a bipolar axis-rough/smooth, hard/soft, slippery/sticky, warm/cold etc., each of these is correlated to a particular physical feature^34–38^. For example, rough and smooth are perceived as a function of spatial period, warm and cool as a function of temperature^33^. Although these bipolar axes reflect the broad categories that textures might vary in, naturally occurring textures could vary on a combination of these categories. Each of these axes could be influenced by the linguistic structures; for example, English-speaking subjects reported adjectives in 3 to 4 dimensions, Japanese-speaking participants’ tactile perceptual space was mapped to 6 dimensions^34–39^. These variations make it possible that the subjects’ linguistic experience could heavily influence texture perception. However, unlike color perception, there have been no systematic cross-language studies of texture perception. One study on Japanese participants showed a greater number of dimensions identified than English speakers, which was attributed to the Japanese having a larger number of sound-symbolic words (SSW) to explain the sensory experience of touch^39^.

Sound symbolic words (SSWs) are a class of verbal labels that are a non-arbitrary label of the sensory stimuli, where the words contain cues about a particular feature of the sensory stimuli it describes. In an investigation of SSWs and tactile stimuli, participants were asked to map the sound-symbolic expressions to the stimuli’ tactile features^40^. Japanese speakers use voiced consonants to express the roughness of materials and voiceless consonants for its smoothness, used bilabial plosive (/p/ and /b/) and voiced alveolar nasal (/n/) consonants for soft, sticky and wet feelings, while that of voiceless alveolar affricate (/t/) and voiceless velar plosive (/k/) consonants for hard, slippery and dry feelings. Another study of Spanish-speaking children shows an association between rough textures and pseudowords with fricatives and smooth textures and pseudowords without fricatives^41^. These studies indicate a strong interaction between perception and linguistic labels in the tactile perceptual space. Thus we set to test the hypothesis that consistent verbal labels increase texture discrimination ability when paired implicitly with a pseudoword verbal label.

The second question we explored in this study is whether the verbal labels’ grammatical properties exert different effects on perception. On the one hand, pseudoword labels suggesting the mere presence of any redundant verbal label can facilitate categorical perception^10,11,13,42^, but others claim that verbal labels’ effect depends on verbal labels’ properties^43,44^. For example, the semantic associations of the verbal label are known to interfere with perception. In the tactile realm, tactile detection improved when the stimulus was preceded by a tactile verb, indicating an interaction between semantic properties of verbal tags and somatosensory processing^43^. Graham et al.^44^ compared the categorization abilities of children after a training phase in which the stimuli were presented along with nouns, adjectives, or stickers. They found that while adjectives helped categorize two stimuli based on similarity, nouns facilitated category formation and generalization. In most languages, individuals interpret tactile stimuli using words that are descriptive or source-based^30^. Extending these categories, we can assume that descriptive responses (for example, smooth) correspond to adjectives while source-based responses (for example, fur) can be mapped to nouns. We manipulated the grammatical identity of the pseudoword verbal labels used in our study such that the pseudowords matched the structure of a noun or adjective in English.

Our study compared performance on a texture discrimination task before and after a training phase where the textures co-occurred with a pseudoword. We methodically co-presented unfamiliar pseudowords with textures that were difficult to discriminate in the congruent condition. In the incongruent condition, we variably co-presented the stimuli such that no consistent association can take place. This procedure was conducted for pseudowords in the grammatical categories of nouns and adjectives. We found that verbal labels facilitated tactile discrimination performance only for the experimental condition after training; however, we did not find any effect of the grammatical category on discrimination performance.

## Results

To evaluate the effect of verbal labels on tactile discrimination ability, we compared the participants’ discrimination sensitivity values (d’ values) in texture discrimination task before and after the training sessions (see methods for more details). We entered these discrimination sensitivity (d′) values in a 2×2 repeated measures ANOVA with training (pre vs post-training) and condition (congruent vs incongruent) as the factors. We found a significant interaction effect (F (1,14) = 5.438, p = 0.035) between the two factors. We also found a significant main effect for training (F (1,14) = 4.892, p = 0.044, d = -0.571), but not for the condition (F (1,14) = 1.206, p = 0.291, d = 0.291).

The paired-samples two-tailed t-test of d′ values between pre and post-training phases showed a significant difference only for congruent condition (t (14) = -3.147, p = 0.007, d = -0.812) but not for the incongruent condition (t (14) = -0.802, p = 0.436, d = -0.207) (Figure 1).Additionally, the change in discrimination sensitivity values (Δd′) after the training phase was significantly greater for the congruent condition than the incongruent condition (t (1,14) = 2.332, p = 0.035, d = 0.602; paired-samples two-tailed t-test) (Figure 2).

**Figure 1:**
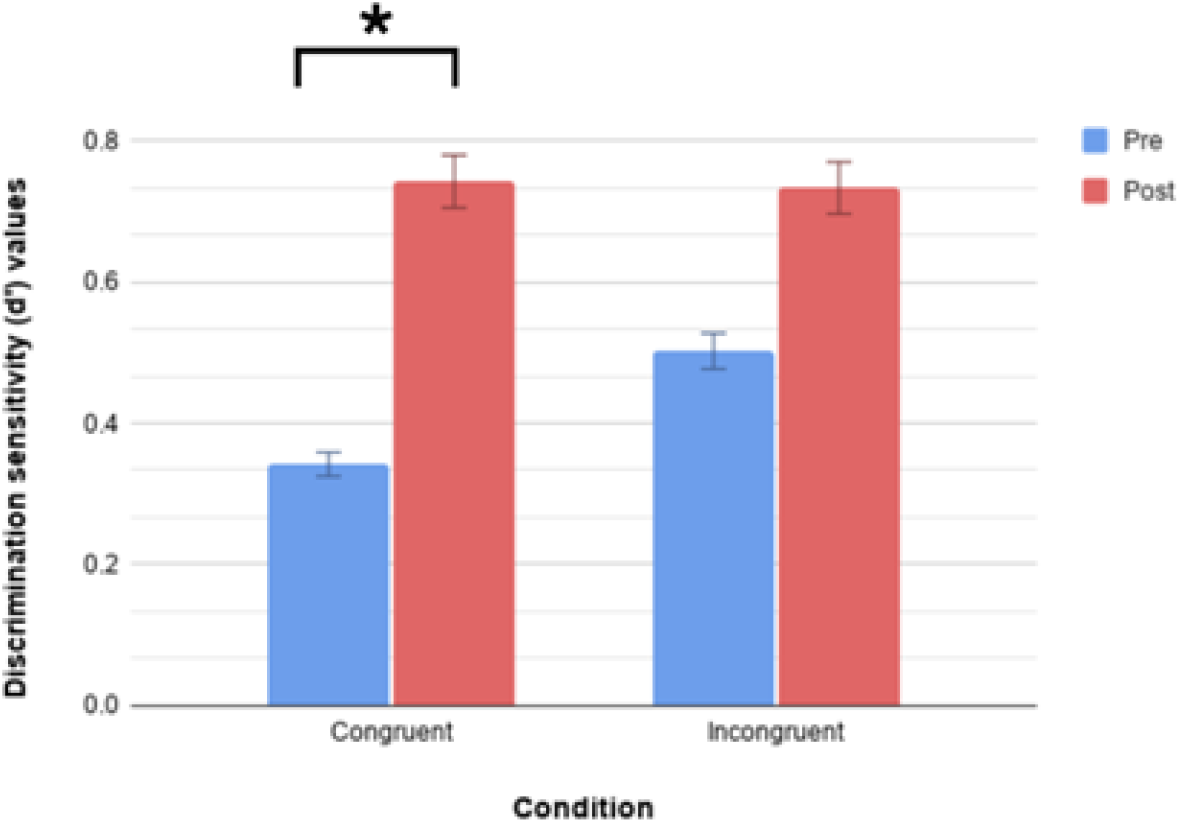
Effect of pseudowords on tactile discrimination. A comparison of the average discrimination sensitivity (d’) values pre- and post-training for 15 participants for the congruent and the incongruent conditions. Discrimination sensitivity significantly improved after training for the congruent condition but not for the incongruent condition. Data shows mean discrimination sensitivity values for 15 participants and the error bars reflect standard error of mean.

**Figure 2:**
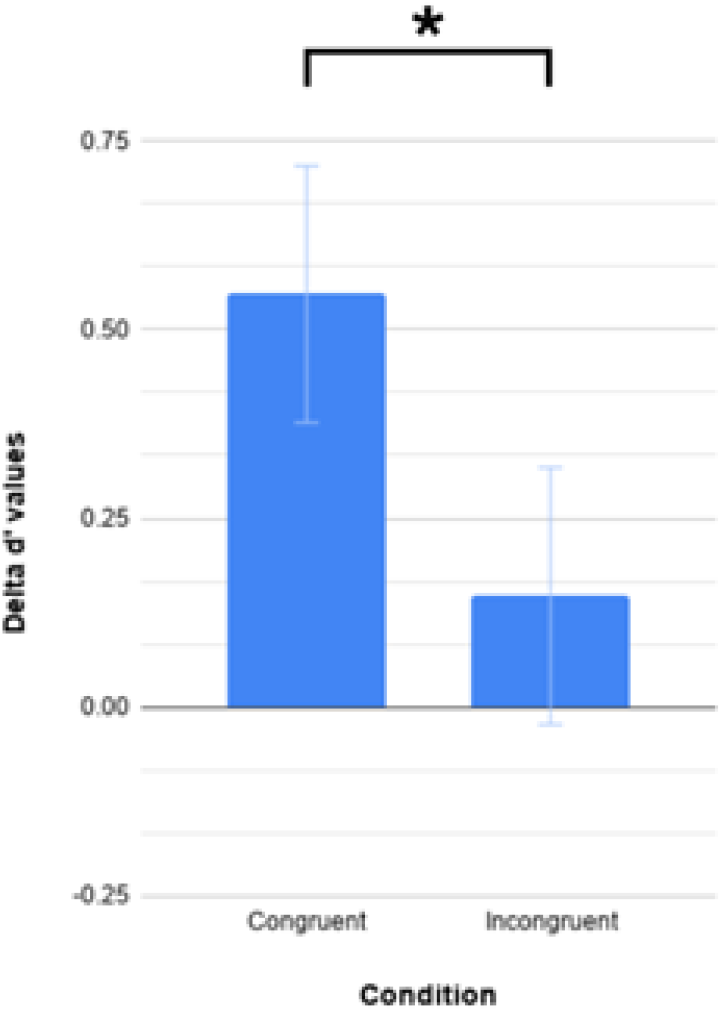
Change in discrimination sensitivity of the congruent and incongruent conditions after training. The difference in the discrimination sensitivity values between the pre- and post-training sessions (Δd’) were significantly different for the congruent and the incongruent conditions. Data shows mean values for 15 participants and the error bars reflect standard error of mean.

Next, we wanted to know if there was an effect of the grammatical category (adjective/noun) pseudoword. To assess this, we entered the d’ values of the congruent condition in a 2×2 repeated measures ANOVA with factors as training (pre vs post-training) and grammatical category (nouns vs adjectives). We found a significant main effect of training, (F (1,14) = 11.565, p = 0.004, d = -0.878) but not for the grammatical category, (F (1,14) = 0.548, p = 0.471, d = 0.191) (Figure 3). No interaction effects were found, (F (1,14) = 0.177, p = 0.680). This suggests that the grammatical category of the verbal label doesn’t play a role in facilitating tactile perceptual discrimination.

**Figure 3:**
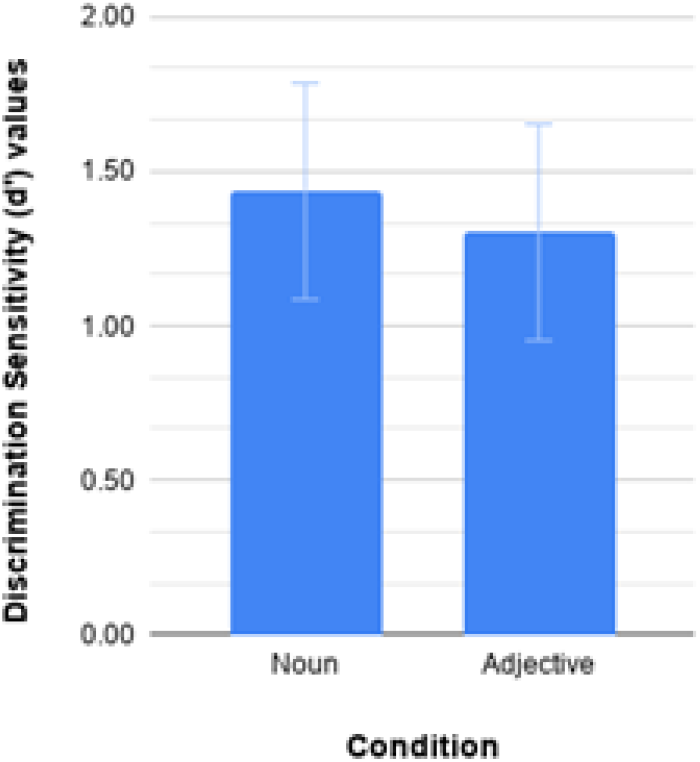
Effect of grammatical category on tactile texture discrimination. A comparison of the average discrimination sensitivity (d’) values pre- and post-training for adjectives and nouns presented during the congruent condition. Discrimination sensitivity significantly improved after training irrespective of the grammatical category of words. There was no statistical difference between the grammatical categories. Data shows mean discrimination sensitivity values for 15 participants and the error bars reflect standard error of mean.

We compared the response bias between the pre-test and the post-test. Using a 2×2 ANOVA, we did not find any main effect of training (F(1,14) = 0.641, p = 0.437) or condition (F(1,14) = 0.138, p = 0.716) as well as any interaction effect (F(1,14) = 2.617, p = 0.128). We also did not find any significant difference in the response bias (t (1,14)=-1.179, p=0.258) between the two conditions in the pre-test.

## Discussion

In the present study, we show that performance in a texture discrimination task improves following a five-day training period where the textures consistently co-occurred with pseudoword verbal labels. The grammatical categories of the pseudowords, i.e, nouns or adjectives, did not affect the discrimination ability. Our findings are in line with previous studies showing that verbal labels facilitate performance on discrimination tasks in other sensory modalities and in the somatosensory system^4,10,28,29^.

In our experiment, we use a within-subject design similar to Miller et al.^28^ that avoids the pitfalls of cross-linguistic studies, which have several confounds such as cultural, social differences and the variation in exposure to the technology used for experiments. In the previous study by Miller et al.^45^, they showed that pseudoword verbal labels improved tactile perception using a 16-pin actuator that delivered patterns of tactile stimuli passively on immobilized fingertips. The pins were spaced 2.5 mm apart and pressed against the skin to create braille-like stimuli. Such stimuli are rarely encountered in nature, so we decided to use stimuli that mimic the everyday tactile experience. Our study differs from the previous studies by using natural textures and unconstrained natural touch where the participants freely moved their fingertip over the textures to perform their task.

The unrestricted use of finger movements shows that the effect of verbal labels on tactile discrimination ability is impervious to participants’ different motor strategies^46–49^. Active and passive touch are considered as fundamentally different paradigms in tactile perception^46,48,50^. Many differences have been shown between these two paradigms of touch; for example, fine textures can only be detected using active touch^51^. Active touch also involves a greater level of top-down influence as individuals adjust the speed, pressure, or orientation of motor movements in response to the sensory feedback^48,51^. So, our study when taken in conjunction with Miller et al.^28^, shows that verbal labels apply to both forms of processing-active and passive touch. We also found an increase in the discrimination ability values with only training, unlike Miller et al.^28^, which could be attributed to the familiar stimuli participants encountered. Similar increases in performance are observed in perceptual learning tasks with tactile stimuli^52,53^.

The verbal labels presented in both the congruent and incongruent conditions varied only in the probability of co-occurrence with the tactile stimuli. In the congruent condition, a pseudoword co-occurred with a particular texture in 70% of the trial, but only in 25% of the trials in the incongruent condition. This ensures that the cognitive load in both conditions are matched. To ensure that participants were paying attention to the pseudowords and the textures, we asked them to report if any pseudowords or textures were repeated after every 10 trials. Furthermore, using pseudowords also prevented the influence of any existing semantic knowledge or conceptual framework. So, we claim that the increase in the discrimination ability is because of verbal labels only.

Even though our results indicate improvement in the tactile discrimination for the congruent condition, there was a difference in the discrimination sensitivity between the two conditions before training. This discrepancy could not have arisen from the inherent differences between the textures used as we counterbalanced the textures across participants. A plausible explanation could be the natural variation in texture perception ability or the motor variations in the active touch strategy, both of which we did not measure^47,48^. Also, the speed of finger movement, the force applied during touch, and the skin area in contact with the surface were not controlled^51^.

This association of verbal labels and sensory stimuli can occur both implicitly and explicitly. Lupyan et al (2007, 2015)^10,13^ reported an improvement in categorization when the participants were instructed to pay attention to the verbal labels corresponding to the visual stimuli. In another study, participants were not explicitly instructed to pay attention^28^. Our verbal label cues were implicitly associated with the textures during the training phase and were not presented during the testing phase. We also show retention of this learning for 24 hours after the last learning phase. Unlike Lupyan et al.’s^10,13^ experiments where the testing phase immediately followed the training phase, the retention of learning of texture-word associations in our study could be compared with more recent experiments that conducted training sessions for 4-5 consecutive days and tested on the following day. However, further studies need to be conducted to study the extent of this effect.

How verbal labels affect perception can be explained through categorical perception, which is the tendency to perceive continuous sensory stimuli as different percepts, like the different colors of the rainbow^8,54–56^. Generally, categorization leads to easy discrimination of stimuli across categories and harder discrimination within a category^56^. This category membership can arise from a shared sensory feature only available to that category^8,9,56^. It is argued that the verbal labels can also act as a shared cue for the sensory stimuli in a category^42^. In our congruent group, where a verbal label was consistently paired with a texture, it acted as an additional and reliable cue for that stimulus, which could have facilitated discrimination. In the incongruent condition, the verbal labels did not co-occur reliably with the textures, so they could not act as an additional cue, so they were all in the same category, making discrimination between the textures difficult.

Categorical perception could change the sensory representations or facilitated attentional mechanisms to enhance discrimination, and our study does not distinguish between the two theories^29,42,55,56^. When categorization occurs, the representation of the stimuli changes, referred to as ‘perceptual warping’ ^42,54^. Here, after the participants learn the verbal label-texture pair, the representations of the textures in the congruent condition are warped into non-overlapping ones as they have unique verbal labels associated with them. This would improve the discrimination performance in the post-test. In the control condition, since there is no unique verbal label, the textures are represented in overlapping representations in similar categories, which leads to poor performance in the test task. An alternate explanation is that paired verbal labels focus attentional mechanisms on the textures’ differences and facilitate discrimination^29,56^.

In our study, the verbal labels’ grammatical category did not show any difference in their effect on tactile discrimination ability. Previous findings report that the grammatical category indicated by the phonetic properties of the label can influence categorization^40,44^. In the English language, nouns and adjectives are supposed to function differently in the categorization process^44^. Nouns represent category names and facilitate better categorization across a category boundary. Adjectives are mostly used to indicate sensory attributes within a category. Since nouns are better suited as verbal labels to discriminate stimuli between categories, we expected nouns to improve the discrimination ability better. However, we did not find this effect. Consistent verbal labels did not differ in their enhancement of tactile discrimination ability depending on the verbal label’s grammatical category. One of the possible reasons for not getting this effect could be that our participants had different first languages even though they were fluent English speakers and writers. Although we tested that individuals could distinguish between nouns and adjectives in the pilot experiment, we did not test it for the participants in the main experiment. Another plausible reason for not getting this effect could be that we selected pseudowords based on the pilot experiment in which participants read the words and categorized them as nouns or adjectives. But in our main experiment, we presented the audio recordings of pseudowords.

In conclusion, our study shows that pseudoword verbal labels improve texture discrimination by facilitating perceptual learning in the tactile modality. Our study adds to evidence for the Linguistic Relativity hypothesis from the somatosensory modality, along with visual, auditory, and olfactory studies. Further studies are needed to understand how the verbal labels influence perception from multiple modalities. Here, information from the somatosensory system can help to understand the nature of this interaction; whether it is at a conceptual, low, mid-level sensory processing or is an attentional process. There is evidence to show differences in neural processing is influenced by linguistic labels are at the earliest sensory processing in the visual cortex^14,16–18^. This idea is disputed by those who say that the effect of language is superficial and that it does not change the way the brain is structured^3,57^. Additional studies of neural sensory representations modified by verbal labels are needed in multiple modalities to show how language affects perceptual processes.

### Materials and Methods

The procedure followed in the present study was approved by the Institute Ethics Committee of Indian Institute of Technology Gandhinagar.

### Experimental Details

In our study, we adopted a within-subject pre-test/post-test design with a five-day long training period in between. Each session was conducted on separate, consecutive days and lasted for about an hour. After taking the informed consent, we explained the procedure of the sessions verbally and a few practice trials were given before the pre-test and first training session. The participants were instructed to touch the textures only using the forefinger of their right hand.

In the pre- and post-test, the participants had to perform a two-alternative forced choice texture discrimination task. In this task, they were asked to actively touch a pair of textures in each trial and report whether they were the same or different. In each trial, participants touched two textures for 1200 milliseconds each, presented one after the other with a variable inter-stimulus interval of 1-3 seconds. An LED light was switched on to indicate when the participants could touch the texture. After this, a buzzer sounded for 300 milliseconds to prompt the participant to report whether the textures were same or different by pressing the red or the blue button respectively on a touchpad. The next trial began as soon as the participants’ response was recorded or 4 seconds after the buzzer sound if no response was made within that duration. Each testing session had 4 blocks of 40 trials and took almost an hour to complete. Participants were given a short break after each block. All possible combinations of textures were presented in a random order in each session.

During the training sessions, each texture was co-presented with a pseudoword. Each condition consisted of 4 distinct textures and pseudowords (2 nouns and 2 adjectives). Here, we exposed the participants to two conditions: the congruent (experimental) condition and the incongruent (control) condition. In the congruent condition, the textures co-occurred with a pseudoword in 70% of the trials and co-occurred with the remaining 3 pseudo-words for 30% of the trials. In the incongruent condition, each texture was paired with each pseudoword belonging to that condition in 25% trials (See Figure 4). Within each block, texture-pseudoword pairings were presented from both the congruent and the incongruent conditions in a random order. The number of texture-pseudoword pairings were constant across all participants and training sessions.

**Figure 4:**
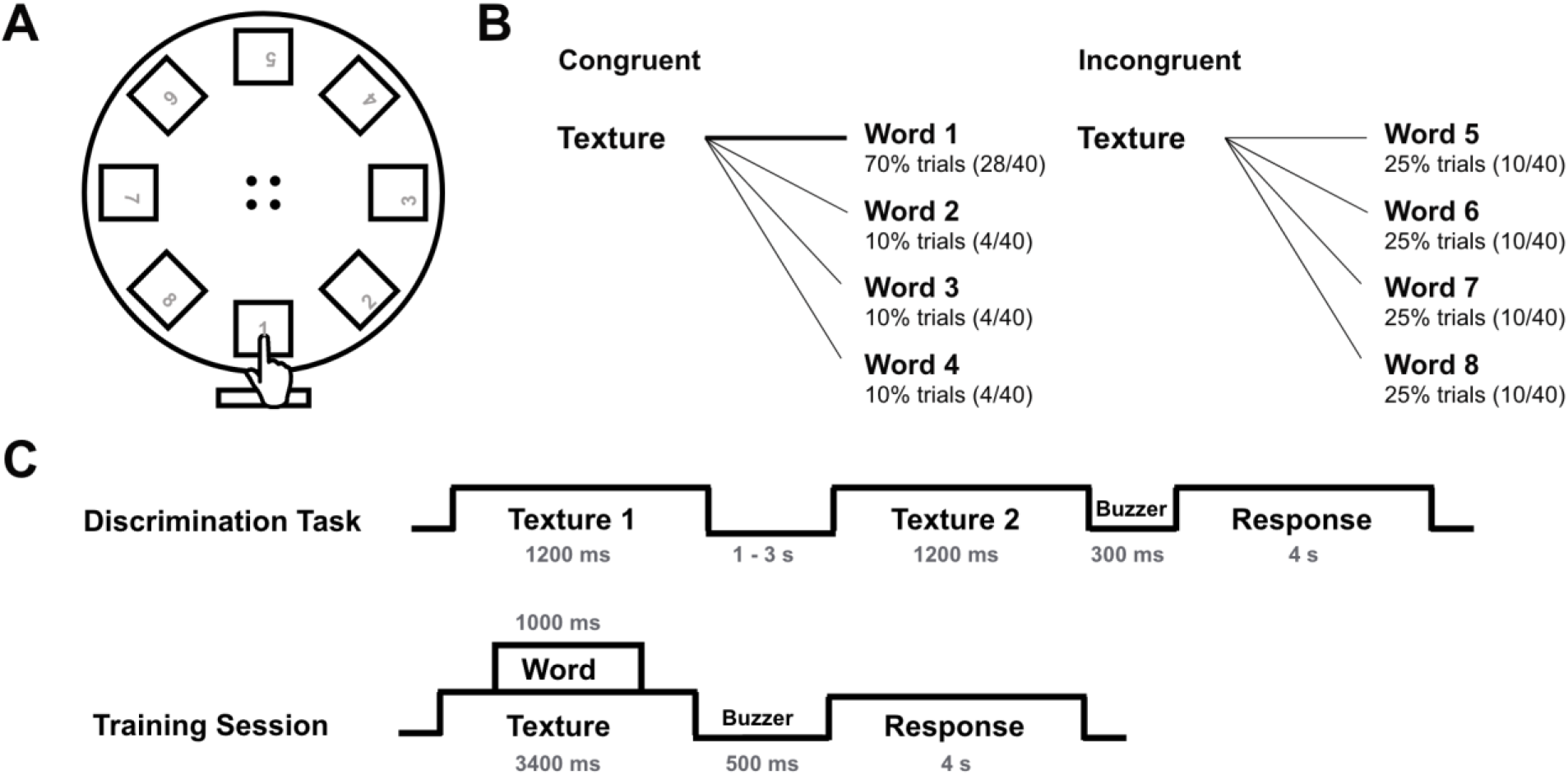
Experimental Setup and Task Design. **(A)** Schematic of the rotating disk apparatus with 8 textures presented in the squares, which were hidden from participants’ vision; **(B)** Textures in the congruent trials were consistently paired with one pseudo-word in 70% trials and variably in the other trials while those in the incongruent condition co-occurred with a pseudowords in 25% trials; **(C)** Trial structure of the texture discrimination task and the training session.

In each trial of the training session, the participant actively touched the texture when the LED light was on for 3400 milliseconds. During this period, an audio clip of the pseudoword was played through the earphones, lasting for 1000 milliseconds. To ensure that the participants paid attention to the stimuli, we asked them to report the number of textures or pseudowords being repeated after every 10 trials, alternating across blocks. They responded by pressing a number (1-9) on the keypad. Each session had 8 blocks of 40 trials, and lasted for approximately one hour. Participants were given short breaks after every two blocks. To eliminate the confounds arising from the differences between groups of stimuli, we counterbalanced the textures across participants in this study, such that one group of textures was presented in the congruent condition for some of the participants and in incongruent conditions for the remaining participants.

## Data Analysis

We analysed the data for the pre-test and post-test to test our hypotheses. We computed the hit rate and the false alarm rate for each participant for both the conditions. The hit rate was computed as the mean correct responses when the pair of textures were different. The false alarm rates were computed as the mean correct responses when the texture pairs consisted of the same textures and were incorrectly perceived as different. These raw scores were converted into standardised z-scores. Then, we computed the discrimination sensitivity values and the response bias according to the Signal Detection theory^58^ using the following formula

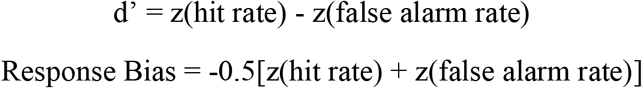

Inferential statistical tests including repeated measures ANOVA and paired-samples t-tests were used to compare the values from the pre-test and the post-test. All statistical analysis was performed using JASP Version 0.14.1 (University of Amsterdam).

### Stimuli and Experimental Setup

#### Experimental Setup

The stimuli were presented to the participants using a custom-made device for the purpose of the current experiment. Textured papers were pasted on 3×3 cm square acrylic pieces placed in slots of the same size on a 5 mm thick acrylic disk of 25 cm diameter precisely cut using a laser machine. This disk was supported on NEMA17 high torque stepper motor (3.1 kg cm, 200 rps, 1.8 degree step angle) firmly upheld on an MDF board base. The motor was controlled using a 5-Ampere dual motor driver compatible with and connected to the Arduino Mega 2560 board and a 12-volt power supply. The verbal stimuli were presented using Philips SHE1350 In-Ear earphones, connected to the Arduino board that also controlled the lighting of the LED bulb when the motor stopped, the ringing of a piezo buzzer to indicate a response. The participant made a response by pressing a key on a 4×4 matrix-membrane keypad that was stored in a 16GB SanDisk MicroSD card connected to the Arduino board via a compatible SD card reader module.

To block the textures from participants’ vision, the entire disk was covered by a 4-side enclosure. Similar to the disk, the box was made using a 5 mm thick acrylic sheet, precisely cut using the laser cutting machine. It had a 3×5 cm slit to allow participants touch the textures on the disk using only one finger. A rectangular piece of foam was kept in front of this enclosure to provide support to participants’ hands. Custom programs were written in Arduino CC version 1.8.10 for two parts of the experiment: texture discrimination task and the training session.

#### Tactile Stimuli

The stimuli consisted of 8 white-colored textured papers of 100 or 120 grams thickness (Conqueror and Rives manufactured by Arjowiggins Creative Papers Ltd). To ensure that these textures are difficult to discriminate in order to avoid any ceiling effects, we conducted a pilot experiment, where 6 participants (4 males, 2 females, mean age = 24.25 years) reported whether a pair of textures was the same or different. All possible combinations of textured papers were presented to each participant. We computed the percentage of correct responses and constructed a matrix with the accuracy for unique texture pairs (See Supplementary Table 1). The results showed that accuracy was low for most texture pairs. Using these results, we divided the textures into two groups for the control and the experimental conditions. There was no overlap between the textures within and between the sets.

#### Verbal Stimuli

Pseudo-words for this study were adapted from Saffran and Theissen’s study^59^ in which they showed infants repeatedly extracted phonological patterns from words. The words were phonologically mapped to two grammatical categories, namely nouns and adjectives. Each pseudo-word had unique phonemes and a constant word length. Since roots and morphemes are independently stored and accessed in the brain^60,61^, we manipulated 3-letter pseudo-words by adding suffixes similar to adjectives (for example, balish, gikous) in English, without allowing any other semantic association. Nouns followed the structure CVCCVC (for example, kupter, pudrat). From the entire pool of pseudo-words constructed for this study, 8 words were selected based on the pilot experiment and categorised into two sets.

In the pilot study, we presented 7 participants (2 females, mean age = 24.14 years) with 96 English pseudo-words mapped to the phonological properties of three grammatical categories (nouns, adjectives, and verbs) by adding an appropriate morpheme. Each pseudo-word was presented on the screen and the participant was asked to categorise it into one of the three grammatical categories using a keypress. We analysed how consistently each word was categorised into a particular grammatical category by the participants (See Supplementary Table 2). Based on these results, we selected 8 pseudo-words that were most frequently perceived to be nouns and adjectives; and randomly divided them into two groups. Each set consisted of two nouns and two adjectives. The audio recordings of these pseudo-words were obtained using text to speech converter (https://soundoftext.com) in a female British English speaker’s voice.

### Participants

Since previous experiments have shown sex differences in haptic texture perception such that men are significantly better than women in the categorisation of tactile stimuli including textures and shapes^62,63^, we recruited only male participants in the present study. 15 male students (mean age = 20.83 years) pursuing graduate or undergraduate degree programs at Indian Institute of Technology Gandhinagar participated in this experiment. They were right-handed, screened using the Edinburgh Handedness Inventory^64^, with fluent English, normal or corrected-to-normal vision, and no calloused fingers. All participants signed an informed consent form before each session of the experiment and were given a monetary compensation for their time.

## Data Availability

The experimental data and the code for the experimental setup is available from the authors upon request.

## Acknowledgements

We would like to thank Jainam Shah, Dr. Madhu Vadali for their help in fabricating the experimental apparatus and Dr. Nishant Choksi for discussion on the linguistic stimuli.

## Author Contributions

IA and LL were involved in conception and design of the study, IA acquired data, IA and LL analyzed and interpreted the data, IA and LL wrote and edited the manuscript.

## Supplementary Tables

**Supplementary table 1:** Percentage of correct responses for unique pairs of textures. To select the textures for the experiment that are difficult to discriminate, we asked 6 participants (2 females, 4 males, mean age = 24.25 years) to discriminate pairs of 8 textured papers. Each participant completed 260 trials with all possible combinations of textures presented in a random order. The table shows the percentage of correct responses in the discrimination task for each unique combination of texture. The discrimination accuracy for most combinations is below or close to 50 percent, showing that the participants could not easily discriminate between them.

| Accuracy | 1 | 2 | 3 | 4 | 5 | 6 | 7 | 8 |
| --- | --- | --- | --- | --- | --- | --- | --- | --- |
| 1 | 62.79 | 25 | 23.26 | 28.26 | 31.82 | 26.09 | 55 | 40.91 |
| 2 |  | 76.09 | 54.55 | 39.53 | 54.55 | 38.1 | 47.62 | 31.82 |
| 3 |  |  | 72.73 | 22.22 | 20 | 28.57 | 40 | 43.48 |
| 4 |  |  |  | 71.43 | 35.00 | 31.82 | 45.45 | 25.00 |
| 5 |  |  |  |  | 63.04 | 31.11 | 42.55 | 39.02 |
| 6 |  |  |  |  |  | 81.40 | 44.19 | 25.00 |
| 7 |  |  |  |  |  |  | 71.78 | 57.78 |
| 8 |  |  |  |  |  |  |  | 63.64 |

**Supplementary table 2:**
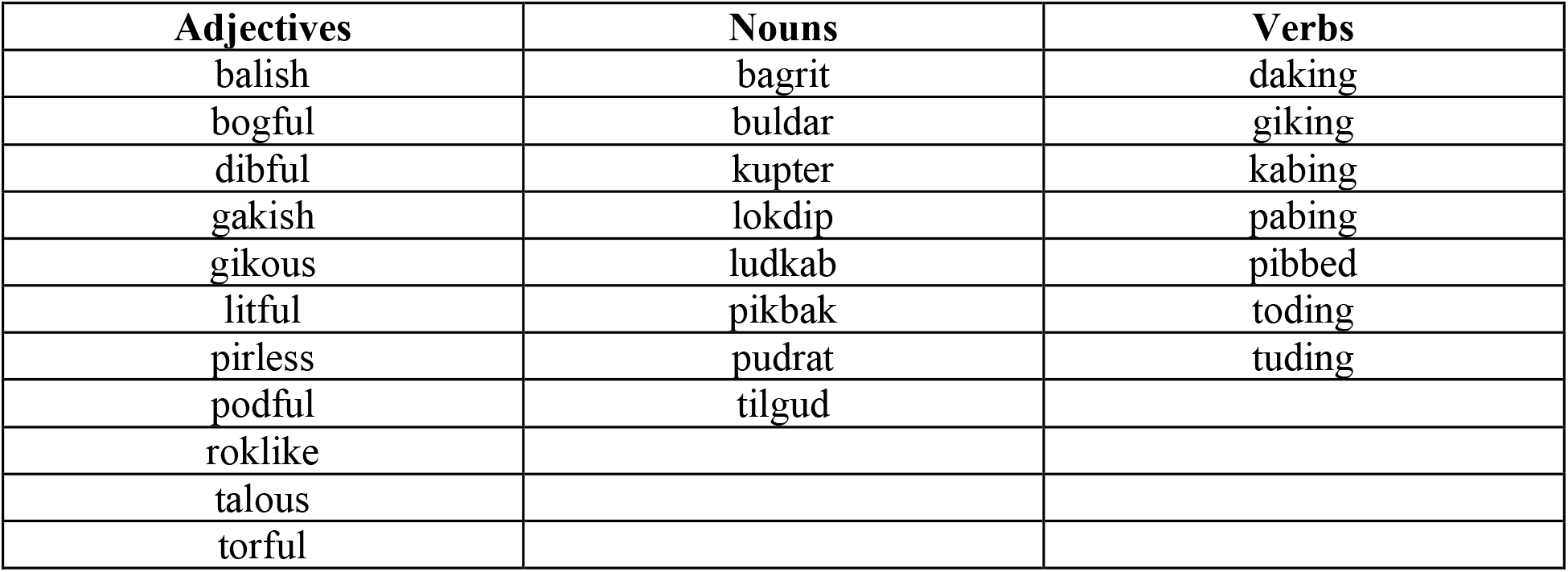
List of words consistently categorised as adjectives, nouns, or verbs by maximum number of participants. To select the pseudo-words for training sessions, we asked 7 participants (2 females, 5 males, mean age = 24.14 years) to categorise pseudowords as nouns, adjectives, or verbs. 32 words per category were randomly presented to the participants. The words that were consistently categorised into three grammatical categories by most participants were shortlisted for the experiment. We compared the average reaction times across 7 participants taken to categorise these pseudo-words. We then selected 4 nouns and 4 adjectives for the experiment.

